# *Ex vivo* functional characterization of mouse olfactory bulb projection neurons reveals a heterogenous continuum

**DOI:** 10.1101/2024.07.17.603915

**Authors:** Sana Gadiwalla, Chloé Guillaume, Li Huang, Samuel JB White, Nihal Basha, Pétur Henry Petersen, Elisa Galliano

## Abstract

Mitral and tufted cells in the olfactory bulb (OB) act as an input convergence hub and transmit information to higher olfactory areas. Since first characterized, they have been classed as distinct projection neurons based on size and location: laminarly-arranged mitral cells with a diameter larger than 20µm in the mitral layer (ML), and smaller tufted cells spread across both the ML and external plexiform layer (EPL). Recent *in vivo* work has shown that these neurons encode complementary olfactory information, akin to parallel channels in other sensory systems. Yet, many *ex vivo* studies still collapse them into a single class, mitral/tufted, when describing their physiological properties and impact on circuit function. Using immunohistochemistry and whole-cell patch clamp electrophysiology in fixed or acute slices from adult mice, we attempted to align *in viv* o and *ex vivo* data and test a soma size-based classifier of OB projection neurons using passive and intrinsic firing properties. We found that there is no clear separation between cell types based on passive or active properties. Rather, there is a heterogeneous continuum with three loosely clustered subgroups: EPL tufted cells, and putative tufted or putative mitral cells in the ML. These findings illustrate the large functional heterogeneity present within the OB projection neurons and complement existing literature highlighting how heterogeneity in sensory systems is preponderant and possibly used in the OB to decode complex olfactory information.

## Introduction

To guide behaviour, concurrent and complex information from the environment must be efficiently decoded and processed by parallel pathways. First described for nociception and vision, parallel processing has been shown to be a hallmark strategy of the brain in sensory systems (Gasser and Erlanger, 1929; Hubel and Wiesel, 1959; Nassi and Callaway, 2009). In the mammalian olfactory system, parallel processing is implemented via two classes of output neurons, mitral and tufted cells, which bring odour information transduced in the olfactory epithelium and pre-processed in the olfactory bulb to higher olfactory areas. While often lumped together and collectively referred to as mitral/tufted cells (M/TCs), recent work has started to highlight that these two classes of projection neurons are rather different.

Mitral cells (MC) are laminarly arranged in the mitral layer (ML) and are known to have the largest somatic diameter (*>*20µm) in the main olfactory bulb (MOB) (Nagayama et al., 2014; Imamura et al., 2020). Conversely, tufted cells (TCs) are diffusely located throughout the external plexiform layer (EPL). TCs are further subdivided into groups based on the their soma position: superficial, middle, internal/deep, from closest to furthest from the glomerular layer (GL), respectively (Schneider and Macrides, 1978; Orona et al., 1984; Nagayama et al., 2014). It should be noted that internal TCs are sometimes called “displaced mitral cells” due to their proximity to the ML (Mori et al., 1983; Ma et al., 2013). TC soma diameter ranges between 10-20µm and is not correlated to their EPL location (Pinching and Powell, 1971; Fukunaga et al., 2012; Nagayama et al., 2014).

Beyond soma size, MCs and TCs are very similar morphologically, both having a primary apical dendrite which ends with a tuft into a single glomerulus (Pinching and Powell, 1971). However, they are differentially connected within the OB network, and superficial TCs receive stronger excitatory inputs from OSNs than MCs (Jones et al., 2020), as well as inhibitory drive from interneurons in the glomerular and granular layer (Christie et al., 2001; Geramita et al., 2016; Liu et al., 2019). Additionally, in higher olfactory areas MCs reach wider territories of piriform cortex (Nagayama, 2010; Igarashi et al., 2012), a difference recapitulated at the molecular level by a differential expression of axon guidance genes (Zeppilli et al., 2021).

Functionally, *in vivo* studies have shown that MCs and TCs encode complementary information thanks to different biophysical characteristics (Balu et al., 2004; Nagayama et al., 2004; Padmanabhan and Urban, 2010; Angelo and Margrie, 2011; Burton and Urban, 2014; Cavarretta et al., 2018; Ackels et al., 2020). Specifically, MCs encode odour concentration while TCs play a key role in odour discrimination (Fukunaga et al., 2012; Igarashi et al., 2012; Burton and Urban, 2014; Nagayama et al., 2014 - but see Chae et al., 2022). As such, TCs exhibit greater firing rates than MCs and have a shorter latency for odour response (Nagayama et al., 2004; Igarashi et al., 2012; Ackels et al., 2020).

Yet, despite this substantial *in vivo* evidence of diversity, few *ex vivo* studies discriminate between MCs and TCs and instead collapse them into an M/TC group. This lack of specificity between OB projection neurons – somewhat unsurprising given that the generation of specific transgenic mouse lines is very recent (Haddad et al., 2013; Koldaeva et al., 2021) – makes it difficult to align results across studies. A decade ago, Burton and Urban started to fill this gap by showing how internal TCs substantially differ from very large neurons in the ML (putative MCs, pMCs) in pre-weaning mice (Burton and Urban, 2014). This seminal study left three important questions unanswered: (a) whether the widely agreed-upon 20µm-diameter classifier maps onto their data, (b) whether smaller cells in the ML are more similar to EPL TCs or to MCs, and (c) whether their results persist in adult animals given that in the first postnatal weeks both TCs and MCs are still undergoing developmental changes (Yu et al., 2015; Tufo et al., 2022). To answer these questions and to harmonize recent *in vivo* studies with *ex vivo* work performed in mouse lines without a specific fluorescent tag for MCs or TCs, this study aims to uncover whether the 20µm-diameter classifier coupled with the intrinsic physiological features of OB projection neurons are sufficient to unequivocally classify MCs and TCs across all layers.

## Materials and Methods

### Animals

Mice of both sexes, aged between postnatal days (P) 21 and 68 were housed under a 12h light-dark cycle in an environmentally controlled room with *ad libitum* access to food and water. In line with the 3R principles we used both wild type mice (C57Bl/6J; Charles River) and surplus animals from transgenic breedings (DATIREScre (B6.SJL-Slc6a3tm1.1(cre)Bkmn/J, Jax stock 006660) and Ai9 (B6.Cg Gt(ROSA)26Sortm9(CAG-tdTomato)Hze/J; Jax stock 007909)) ongoing in the laboratory. The integrity of the transgenic lines was ensured by generating breeders via back-crossing heterozygous carriers with C57Bl6 animals specifically bought biannually from Charles Rivers. All experiments were performed at the University of Cambridge in accordance with the Animals (Scientific procedures) Act 1986 and with AWERB (Animal Welfare and Ethical Review Board) approval.

### Immunohistochemistry

Mice were anesthetized with an overdose of pentobarbital and then perfused with 20 mL PBS with heparin (20 units.mL^-1^), followed by 20mL of 1% paraformaldehyde (PFA; TAAB Laboratories; in 3% sucrose, 60 mM PIPES, 25 mM HEPES, 5 mM EGTA, and 1 mM MgCl_2_). Olfactory bulbs were dissected and post-fixed in 1% PFA for 2-7 days and embedded in 5% agarose and sectioned into 50µm slices using a vibratome (VT1000S, Leica). Free-floating slices were washed with PBS and incubated in 5% normal goat serum in PBS/Triton/azide (0.25% Triton, 0.02% azide) for 2 hours at room temperature and then incubated in primary antibody solution (in PBS/Triton/azide) for 2 days at 4°C. Primary antibodies and their working dilutions included SMI-32 neurofilament H non-phosphorylated (SMI-32; mouse, Biolegend; 1:1000), Ankyrin-G (Guinea Pig, Synaptic Systems, 1:500),), and tyrosine hydroxylase (rabbit, Sigma Millipore AB152, 1:500). Following primary incubation, slices were washed 3 times in PBS for 5 minutes before secondary antibody solution (species-appropriate, Life Technologies Alexa Fluor-conjugated) 1:1000 in PBS/Triton/azide for 3 hours at room temperature. Slices were then washed in PBS and incubated in 0.2% Sudan black in 70% ethanol at room temperature for 3 minutes to minimize autofluorescence and mounted on glass slides (Menzel-Gläser) with FluorSave (Merck Millipore).

### Fixed-tissue imaging and analysis

Images were acquired with a laser scanning confocal microscope (Carl Zeiss LSM 900) using appropriate excitation and emission filters, a pinhole of 1 AU and a 40x oil immersion objective. Laser power and gain were set to prevent signal saturation in channel images for localisation analysis. All quantitative analysis was performed with Fiji (ImageJ).

For soma size analysis, images were taken with a 1x zoom (0.415µm/pixel), 512 × 512 pixels in z-stacks with 1µm steps. In all animals, images were sampled from the rostral third, middle third and caudal third of the OB. To avoid selection bias, all cells present in the stack and positive for the SMI-32 were measured using Fiji/ImageJ. Soma area, Feret’s diameter (*i*.*e*., the longest distance between any two points), roundness defined as 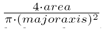, and circulaity defined as 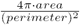, was measured at the single plane including the cell’s maximum diameter by drawing an ROI with the free-hand drawing tool. Cells were classified based on their location within the ML and EPL. If their soma fell clearly within the boundaries of the ML it was determined to be a putative mitral or tufted cell. Alternatively, neurons whose soma was fully located in lower third of the EPL (*i*.*e*. the EPL third closest to the ML) were classed as EPL tufted cells.

For AIS identification, images were taken with 2x zoom, 512 × 512 pixels (0.138µm/pixel) in z-stacks with 0.30µm steps. Laser power and gain settings were adjusted to prevent signal saturation in the AIS label AnkG. The cellular marker SMI-32 signal was usually saturated to enable clear delineation of the axon. Distance from soma and length was measured in Fiji/ImageJ using the View5D plugin which allows for 3D manual tracing of cell processes. The AIS distance from soma was calculated as the neurite path distance between the start of the AIS (proximal part where AnkG staining was clearly identifiable) and the intersection of its primary parent process (the axon). AIS length was calculated by following AnkG staining along the course of the axon from the AIS start position to the point where AnkG staining was no longer identifiable and only SMI-32 straining was present along the axon.

### Electrophysiology

Mice were decapitated under isoflurane anaesthesia. The OB was removed and transferred into ice-cold slicing medium containing the following (in mM): 240 sucrose, 5 KCl, 1.25 Na_2_HPO_4_, 2 MgSO_4_, 1 CaCl_2_, 26 NaHCO_3_ and 10 D-Glucose continuously bubbled with 95% O2 and 5% CO2. Horizontal slices (300µm) of the OB were cut using a vibratome (Campden Instruments 7000SMZ-2 or Leica VT1000S) and stored in ACSF containing the following (in mM): 124 NaCl, 2.5 KCl, 1.25 Na_2_HPO_4_, 5 MgSO_4_, 2 CaCl_2_, 26 NaHCO_3_ and 15 D-Glucose, maintained in 95% O2 and 5% CO2 for at least one hour at room temperature before experiments began. Cells were visualized with an upright microscope (BX51W1; Olympus) using a 40X water immersion objective with visible light camera (quantalux sCMOS; ThorLabs) and classified based on their location in the MCL and EPL. Cells were targeted for patch-clamp if the entirety or over half of their soma lay between the boundaries of the MCL. Internal tufted cells were targeted based on their location in the lower third of the EPL (Mori et al., 1983; Kishi et al., 1984; Shepherd, 2004).

Whole-cell patch-clamp recordings were amplified and digitized using an EPC-9 (HEKA Electronik GmbH) or MultiClamp 700B (Molecular Devices) and Digidata 1550B (Molecular Devices) at physiologically relevant temperatures (30±2°C), maintained with an in-line heater (SH-27B and TC-344C, Warner Instruments). All signals were Bessel filtered at 10kHz, with experiments recorded at 20kHz and single spike recordings filtered at 200kHz. Recordings were excluded if series resistance (evaluated by -10mV voltage steps following each test pulse) was *<*40 M for putative mitral cells with a capacitance *>*40pF, and *<*30 M for putative tufted cells with a capacitance *<*40pF. Protocols were also excluded if these values varied more than 20% over the course of the experiment and holding current values exceeded -300pA. Fast capacitance was compensated in on-cell configuration. Cell capacitance was calculated by measuring the area under the curve of a transient capacitive current induced by a -10mV step after subtracting the steady-state current induced by the voltage pulse. Recording electrodes (30-0094 and 30-0062 Harvard Apparatus) were pulled using a horizontal or vertical puller (P-87, Sutter Instruments; PC-100, Narishige) to achieve a tip resistance of 1.5–3.0 M (larger tips for putative mitral, smaller for putative tufted and EPL tufted) when filled with a potassium-gluconate intracellular solution containing the following (in mM): 124 K-gluconate, 9 KCl, 10 KOH, 4 NaCl, 10 HEPES, 28.5 sucrose, 4 Na_2_ATP, 0.4 Na_3_GTP (pH 7.25-7.35; 290 MOsm) and Alexa-488 (Thermo Fisher Scientific, 1:150).

To precisely measure soma size and ensure fluorophore diffusion throughout the entire soma, cells were patched for at least 10-15 minutes prior to capturing a snapshot of their Alexa Fluor 488-filled soma with LED excitation (LED1B; ThorLabs) using the appropriate excitation and emission filters (ET575/50m; CAIRD Research UK). Quantitative analysis was performed in Fiji/ImageJ. Diameter and area were calculated by drawing a ROI using the free-hand drawing tool. In a subset of recordings, biocytin (Sigma; 2%) was added to evaluate morphology. These slices were fixed with 1% PFA in PIPES overnight and then incubated with 1:1000 streptavidin Alexa Fluor 488 conjugate in PBS/Triton/azide for 2h at room temperature to reveal the biocytin filling.

In current-clamp mode, experiments were only evaluated if their holding voltage was -60 ± 3mV. For action potential (AP) measurements, injections of 10ms current steps of increasing amplitudes were applied until current threshold was met, and the cell fired an action potential (Vmax *>* 0mV). Repetitive firing properties were measured with injections of 500ms current steps starting at 0mV of increasing amplitudes (*△*5-35) until the neuron passed its maximum firing frequency. Sag potentials were evoked by injecting a 500ms current injection starting from -300pA to -700pA. Exported traces were analyzed using either ClampFit (pClamp 10, Molecular Devices) or custom-written scripts in MATLAB (MathWorks)(Galliano et al., 2019).

### Quantification of passive and active electrophysiological properties

Quantification of action potential properties was calculated from the first action potential evoked by the weakest suprathreshold input. Current threshold was defined as the minimum threshold needed to elicit the first action potential. Voltage threshold was taken as the potential at which dV/dt first passed 10V/s and the maximum of depolarisation. Peak was the highest voltage an action potential reached. Action potential amplitude was the difference between the voltage threshold and the action potential peak. Spike width was measured at the midpoint between voltage threshold and maximum voltage. Afterhyperpolarisation values were measured from responses to 500ms current injection from the local voltage minimum after the first spike fired at rheobase.

For repetitive firing properties, input-output curves were created by counting the number of spikes fired at each level of injected current density. The slope of the input-output curve was measured between the first sweep with a non-zero action potential and the sweep where the maximum number of action potentials was fired. The following parameters were measured only from the sweep with the maximum number of action potentials were fired: (a) first action potential delay which measured the time interval between the start of the current injection and the peak of the first action potential; (b) peak of the first action potential; (c) action potential frequency. To measure variability in firing pattern, the coefficient of variance (CV) of the inter-spike interval (ISI) was calculated across current injections and at the sweep that fired the maximum number of action potentials. CV was calculated as the ratio of the standard deviation of ISIs to the mean ISI of the cell. Firing pattern variability measure CV2 was calculated as mean value of (2*abs((ISIn+1 – ISIn))/(ISIn+1 + ISIn)) (Holt et al., 1996). Sag potentials were evaluated as done previously (Angelo and Margrie, 2011) where sag index was calculated as the ratio between the peak (minimum within the first 100ms) and steady state (mean of final 50ms) currents normalised to holding voltage.

### Statistical Analysis and data availability

Statistical analyses were carried out using Prism (GraphPad) or MATLAB (MathWorks). “N” refers to number of animals and “n” indicates number of cells. Normality of sample distribution was assessed with the D’Agostino and Pearson omnibus test and their parametric or non-parametric tests used accordingly. *α* values were set at 0.05, and all comparisons were two-tailed. Post-hoc tests after nested ANOVAs were done using the two-stage linear step-up procedure of Benjamini, Krieger and Yekutieli. K-means clustering of immunohistochemistry and electrophysiology data were executed in MATLAB (MathWorks) using custom written scripts which included an evaluation of silhouette coefficient. Cluster numbers were chosen based on the silhouette coefficient that was closest to one. Principal component analysis (PCA) of electrophysiological data was performed in Prism (GraphPad). Principal components (PCs) were selected based on the Kaiser rule, where PCs were selected only if they had an eigenvalue greater than one. All PCAs were unsupervised. Loading scores were calculated based on standardised data using the following formula: (Eigenvector*sqrt(Eigenvalue)). Post publication, the full dataset will be released on the University of Cambridge Apollo repository under a CC-BY license.

## Results

### Soma size and morphology of mitral and tufted cells is heterogeneous

Two criteria have been traditionally used to classify MCs and TCs: the location and size of their somas. There is historical consensus in the literature that MC somas lie entirely or predominantly in the ML and that they are large, with a longest-axis diameter *>*20µm and a corresponding area *>*350µm^2^. Conversely, TC somas are spread across the EPL and on average smaller than MCs’ – longest-axis diameter *<*20µm, area *<*230µm^2^ (Mori et al., 1983; Orona et al., 1984; Ezeh et al., 1993; Royet et al., 1998; Nakajima et al., 2001; Shepherd, 2004; Nagai et al., 2005; Fukunaga et al., 2012; Igarashi et al., 2012; Nagayama et al., 2014). We first investigated whether this historical diameter boundary could be used as a reliable binary classifier of mitral vs tufted identity in fixed OB tissue. We identified M/TCs with antibodies against the neurofilament marker protein, SMI-32, which labels the cell body, axon and dendrites of bulbar excitatory neurons. To unequivocally define the upper border of the EPL, we co-stained the tissue with antibodies against tyrosine hydroxylase (TH) to label glomerular layer’s dopaminergic neurons (Fig. 1A). Using confocal microscopy we acquired 3D z-stacks where we sampled SMI-32-positive neurons in the lower third of the EPL (*i*.*e*., internal TCs) and in the ML. We found that TCs in the EPL had a mean diameter of 15.73µm and mean area of 115.4µm^2^ (Fig. 1B, green circles). In the ML, however, we found that soma sizes were more heterogeneous – mean diameter 20.61µm, range 9.46-39.79µm; mean area 180.4µm^2^, range 19.47-513.22µm^2^ (Fig. 1B, cyan circles; area-diameter linear regression R^2^=0.74; soma diameter frequency distribution Fig. 1C).

**Figure 1.**
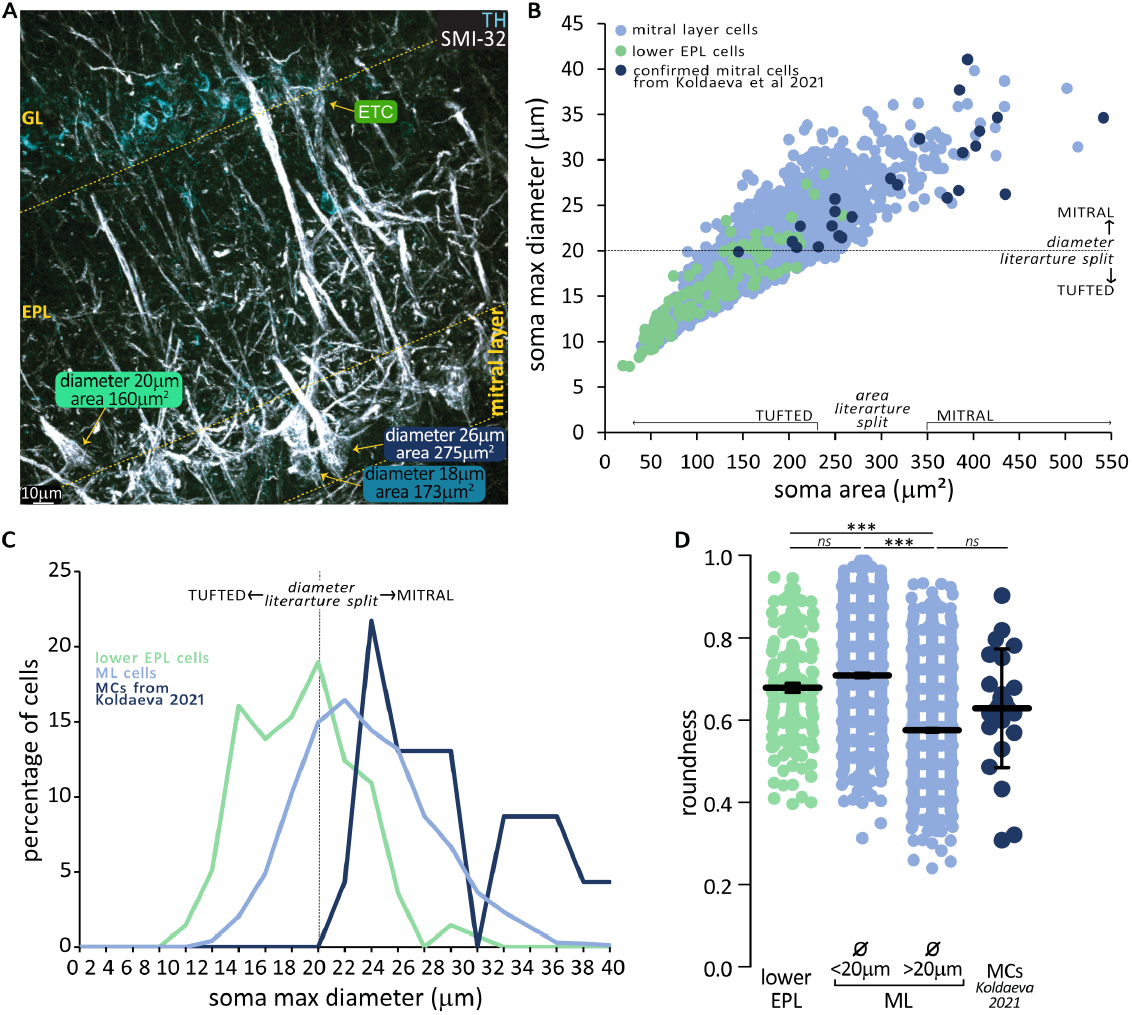
OB projection neurons in the mitral layer include differently sized and shaped cells. **(A)** Example image of the olfactory bulb outer layers. Excitatory projection neurons labelled with antibodies against neurofilament marker protein (SMI-32, white) span the external plexiform layer (EPL) and the mitral layer (ML). Dopaminergic interneurons stained with antibodies against tyrosine hydroxylase (TH, cyan) indicate the location of the glomerular layer (GL), adjacent to which are the SMI-32 positive somas of the excitatory interneurons external tufted cells (ETCs). **(B)** Correlation of soma area and maximum diameter for OB projection neurons located in the lower EPL (green, n=137) or ML (cyan, n=1671). Blue circles represent confirmed mitral cells from the Lbhd2-CreERT2 transgenic mouse line, meta-analysed from Koldaeva et al 2021 (n=22). The canonical diameter and area dividers between mitral and tufted cells are indicated on the axes. **(C)** Frequency distribution of maximum diameters for OB projection neurons located in the lower EPL (green, n=137), ML (cyan, n=1671), and confirmed mitral cells from the Lbhd2-CreERT2 transgenic mouse line **(D)** Soma roundness [(4*area/(π*major_axis^2^)] of EPL tufted cells (green) and confirmed mitral cells (blue) compared to ML cells (cyan) when split by the 20µm max diameter classifier proposed in the literature. Circles are individual cells, lines are mean ± SEM, *p<0.05, **p<0.01, ***p<0.001.

To further describe this heterogeneity, and given that MCs have been further divided according to soma shape (Kikuta et al., 2013), we calculated the soma roundness at its longest-axis diameter. Smaller cells in the ML (diameter *<*20µm) were rounder than ML cells with a diameter *>*20µm, but they did not differ in roundness nor in area from EPL TCs (Kruskal Wallis with Dunn’s corrections, p*<*0.0001; Fig. 1D). Moreover, when we performed a secondary analysis of published data in confirmed MCs labelled in the Lbhd2-CreERT2 transgenic mouse line (Koldaeva et al., 2021; n=22, mean diameter 27.22µm, mean area 317.868µm^2^, mean roundness 0.63; Fig. 1B-D, dark blue dots/line), we found that these neurons were similarly sized and shaped as the largest ML cells (one-way ANOVA nested on mouse or figure with Tukey’s corrections, roundness F(3,22)= 32.95, pMC vs. Koldaeva et al.,2021 p = 0.32; area: F(3,22)= 14.90, pMC vs. Koldaeva et al.,2021 p = 0.05; diameter: F(3,22)=16.56, pMC vs. Koldaeva et al.,2021 p=0.64). In summary, our data is broadly in agreement with the literature in that TCs in the EPL are round and have on average somas smaller than 20µm, and that the ML contains some very large and more ovoidal neurons. However, we found no clean split as advocated in previous studies: first, a minority of large soma neurons are present in the EPL, and second, over a third of ML neurons are small and round. These latter cells are more similar in size and shape to EPL TCs than ML MCs, and we tentatively class as putative ML tufted cells (pTCs).

### Unbiased k-means analysis identifies a capacitance threshold to classify mitral and tufted cells

Are pTCs in the ML not only morphologically but also physiologically like EPL TCs? To evaluate their electrophysiological properties, we employed whole-cell patch clamp in acute slices, where soma location could be easily determined (Fig. 2A). Soma diameter of the patched cells was analysed in light microscopy configuration and subsequently confirmed with LED fluorescence after filling the recorded neuron with an Alexa Fluor 488-supplemented intracellular solution (Fig. 2B). Recognizing the well-known disparity between live and fixed samples due to fixation-induced tissue shrinkage (Boonstra et al., 1983) and given that our earlier analysis failed to reveal a clear separation using the 20µm diameter classifier in fixed tissue (Fig. 1B-C), we decided to adopt an alternative classification strategy. To correlate morphology with electrophysiology and ultimately distinguish between pMCs and pTCs based on soma size-related passive properties, we implemented an unbiased k-means algorithm based on capacitance, a reliable indicator of somatic size. We input the diameter and capacitance of recorded ML cells into the k-means algorithm, which yielded two distinct clusters with centroids at 17.64µm diameter / 29.23pF capacitance (representing pTCs) and 22.51µm diameter / 61.87pF capacitance (representing pMCs). These two clusters could be separated by a capacitance-based classifier of 45pF (Fig. 2C). Using this fully unbiased classifier, we compared mean capacitance of pTCs and pMCs with those of TCs recorded in the EPL (41.63 ± 3.32 pF; max diameter 18.49 ± 0.89 µm) and confirmed that only pMCs are significantly different (Kruskall-Wallis test with Dunn’s corrections eplTCs vs pMCs p*<*0.001; Fig. 2D). In line with this result and in further support of the capacitance classifier, we found that input resistance (Ri, also partially dependent on soma size) was significantly lower in pMCs than in both pTCs and eplTCs (eplTC 205±39 mΩ, pTC 275±50 mΩ, pMC 110±13 mΩ; Kruskall-Wallis test p*<*0.001, Dunn’s corrections eplTCs vs pMCs p*<*0.05, pTCs vs pMCs p*<*0.001; Fig. 2E).

**Figure 2.**
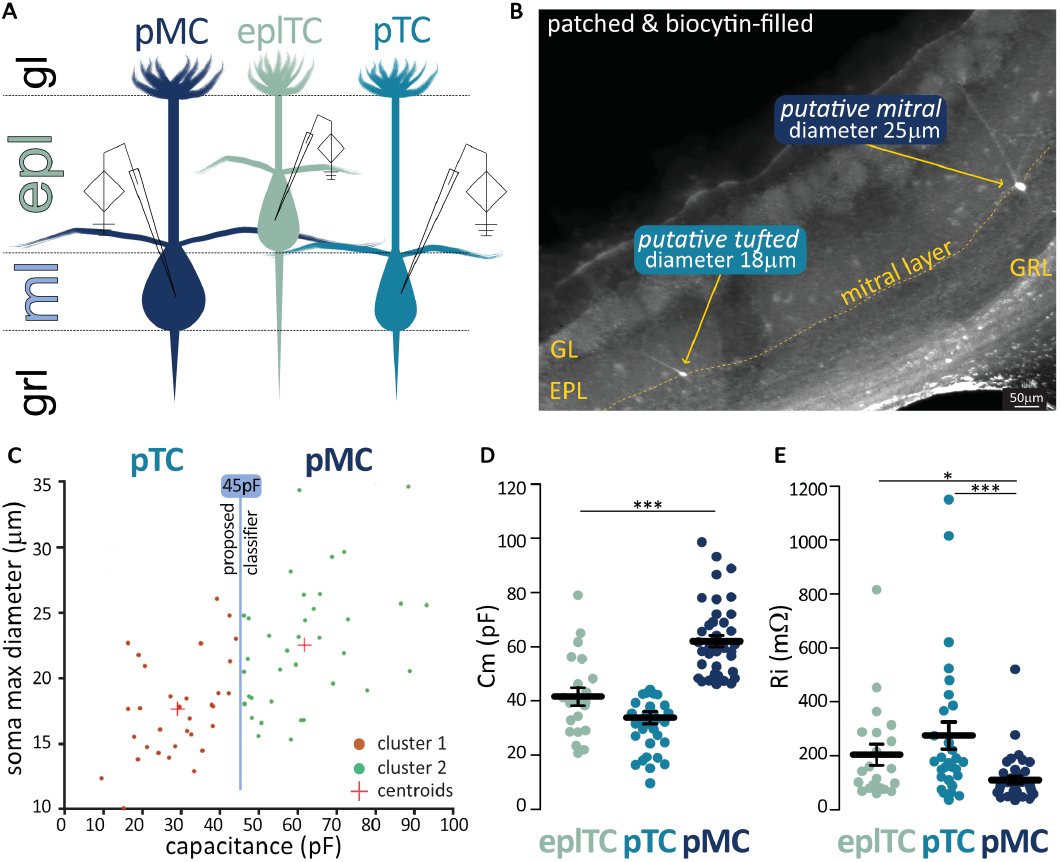
Soma size and passive electrical properties differ between OB projection neurons. **(A)** Schematic representation of the location of OB projection neurons targeted for whole cell patch clamp recordings in acute horizontal brain slices from juvenile mice. **(B)** Two cells in the mitral layer, a putive tufted (pTC) and a putative mitral (pMC) patched with biocytin-supplemented intracellular solution and postfixed for morphological analysis. **(C)** Unbiased k-means analysis of soma size of patched cells in the mitral layer returns two clusters separable by a 45pF capacitance classifier. **(D-E)** Membrane capacitance (Cm) and input resistance (Ri) in epITC (n=21) and mitral layer’s pTC (n=28) and pMC (n=42) classed using the 45pF divider. Circles are individual cells, lines are mean ± SEM, *p<0.05, ***p<0.001. GL = glomerular layer, EPL = external plexiform layer, GRL = granule cell layer.

In summary, we successfully identified a purely electrophysiological measure – capacitance – that can be used in live tissue to attempt a classification of ML cells into the two subgroups.

### Sag voltages are not significantly different between principal neurons in EPL and ML

The presence of a hyperpolarization-activated cation current (I_h_/ sag currents), which are important determinants of a neuron’s intrinsic excitability (Combe and Gasparini, 2021), have been shown to be variable in ML cells (Angelo and Margrie, 2011; Angelo et al., 2012). To assess if this reported heterogeneity mapped on the pTCs/pMCs subclasses, we injected increasing levels of hyperpolarizing current into ML and EPL principal neurons (Fig. 3A-B). We found no differences between pMCs, pTCs and eplTCs in the raw measurements of sag peak and steady state voltage, nor in the combined measure of sag amplitude and index (Angelo and Margrie, 2011) (Fig. 3C-F; ANOVA with Tukey or Kruskall-Wallis with Dunn, all p¿0.2; Table 2). Of note is the variability across all groups, but especially the pTCs (sag amplitude interquartile range and CV: eplTCs = 18.25mV – 97%, pTCs = 4 4.881mV, - 163%, mTCs = 10.48mV – 174%; sag index interquartile range and CV: eplTCs = 0.23 – 17%, pTCs = 0.44, - 44%, mTCs = 0.13 – 21%; Bartlett’s test for equal variances: amplitude p*<*0.001, index p*<*0.01). Across all three groups, approximately half of the cells fire on rebound following the hyperpolarization-induced voltage sag (eplTCs = 50%, pTCs = 58%, pMCs = 48%; Chi-square test, p=0.33).

**Figure 3.**
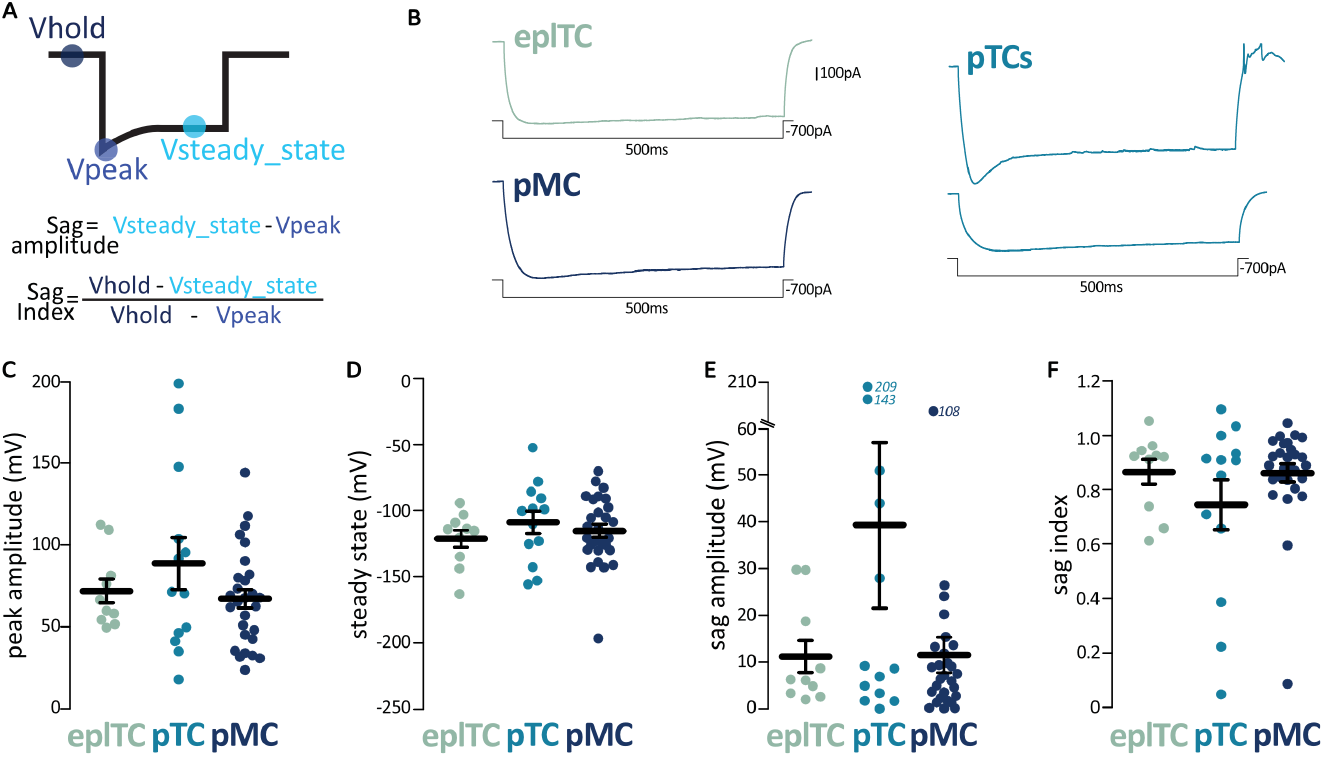
The depolarization response to hyperpolarization (sag potential) is extremely variable among OB projection neurons. **(A)** Schematic visualization of the voltage sag analysis parameters and formulas used to calculate sag amplitude and index. **(B)** Example traces of the voltage sag response to hyperpolarizing current injections in epITC (green, n=10), pTCs (cyan, n=13) and pMC (blue, n=28). Note the variability in pTCs. **(C-F)** Peak amplitude, steady-state voltage, sag amplitude and sag index in the three classes of OB projection neurons. Circles are individual cells, lines are mean ± SEM; further quantification and statistical analysis in Table 1.

Overall, our data confirm the sag variability reported in the literature, but this heterogeneity does not map onto the pTC and pMC subgroups.

**Table 1:** Intrinsic electrophysiological properties of OB projection neurons. Mean values ± SEM of sag currents, action potential properties, and repetitive firing properties for epITCs, and mitral layer’s pTCs and pMC. Statistical differences between cell types were calculated independently with a one-way ANOVA with Tukey’s post hoc correction for normally-distributed data (A), or with a Kruskal-Wallis test with Dunn’s post hoc correction for non-normally distributed data (“KW”). Individual data points and example traces are presented in Figures 3-4-6.

| Intrinsic electrophysiological properties |  |  |  |  |  |  |  |
| --- | --- | --- | --- | --- | --- | --- | --- |
|  | epITC mean<br>± sem, [N] | pTC<br>mean ±<br>sem, [N] | pMC mean<br>± sem, [N] | ANOVA /<br>Kruskal-Wallis<br>p-value | Tukey's / Dunn's<br>multiple comparisons<br>p-value |  |  |
|  |  |  |  |  | EpITC<br>vs pTC | epITC<br>vs<br>pMC | pTC vs<br>pMC |
| SAG POTENTIALS PROPERTIES |  |  |  |  |  |  |  |
| Peak amplitude (mV) | 71.97 ±<br>7.24 [10] | 88.7 ±<br>15.82<br>[13] | 67.27 ± 5.63<br>[28] | A, 0.24<br>(F=1.46) | ns | ns | ns |
| Steady State ( mV) | -121.3 ± 6.5<br>[10] | -109 ±<br>8.39 [13] | -115.5 ±<br>4.89 [28] | KW, 0.52 | ns | ns | ns |
| Sag Amplitude (mV) | 11.20 ±<br>3.43 [10] | 39.28 ±<br>17.80<br>[13] | 11.53 ± 3.80<br>[28] | KW, 0.63 | ns | ns | ns |
| Sag Index | 0.86 ± 0.05<br>[10] | 0.74 ±<br>0.09 [13] | 0.86 ± 0.03<br>[28] | KW, 0.74 | ns | ns | ns |
| ACTION POTENTIALS PROPERTIES |  |  |  |  |  |  |  |
| Current threshold<br>(pA/pF) | 9.63 ± 1.64<br>[13] | 5.46 ±<br>0.87 [13] | 3.06 ± 0.37<br>[27] | KW,<br><0.001 | ns | *** | * |
| Voltage threshold (mV) | -30.22 ±<br>2.18 [13] | -37.26 ±<br>2.76 [13] | -43.14 ±<br>1.14 [27] | A, <0.001<br>(F=13.14) | ns | *** | ns |
| Maximum Voltage (mV) | 10.48 ±<br>2.01 [13] | 14.89 ±<br>2.24 [13] | 14.10 ±<br>1.76 [27] | KW, 0.40 | ns | ns | ns |
| Peak Amplitude (mV) | 40.17 ±<br>1.83 [13] | 58.56 ±<br>4.60 [13] | 58.29 ±<br>2.06 [27] | KW,<br><0.001 | ** | *** | ns |
| Width at Half-Height<br>(ms) | 0.80 ± 0.08<br>[13] | 0.93 ±<br>0.06 [13] | 0.83 ± 0.03<br>[27] | KW, 0.25 | ns | ns | ns |
| Minimum voltage (mV) | -46.22 ±<br>2.05 [15] | -54.03 ±<br>2.30 [11] | -56.26 ±<br>1.87 [27] | A, 0.003<br>(F=6.41) | ns | ** | ns |
| After hyperpolarization<br>amplitude (AHP, mV) | 17.13 ±<br>1.57[15] | 12.61 ±<br>1.18 [11] | 16.59 ±<br>1.10 [27] | A, 0.08<br>(F=2.58) | ns | ns | ns |
| REPETITIVE FIRING PROPERTIES |  |  |  |  |  |  |  |
| Rheobase (pA/pF) | 3.03 ± 0.69<br>[14] | 1.59 ±<br>0.29 [11] | 1.82 ± 0.31<br>[28] | A, 0.08<br>(F=2.58) | ns | ns | ns |
| Slope I/O Curve<br>(Hz/pA/pF) | 7.98 ± 1.38<br>[13] | 6.33 ±<br>1.24 [11] | 7.21 ± 1.01<br>[26] | A, 0.72<br>(F=0.34) | ns | ns | ns |
| First AP Delay (ms) | 109.8 ±<br>20.6 [15] | 169.5 ±<br>42.1 [11] | 161.1 ±<br>29.1 [27] | KW, 0.78 | ns | ns | ns |
| n APs at Rheobase | 8.14 ±<br>2.95 [14] | 5.36 ±<br>1.22 [11] | 8.64 ±<br>2.56 [28] | KW, 0.86 | ns | ns | ns |
| Max Number of APs | 29.87 ±<br>4.76 [15] | 22.27 ±<br>3.65 [11] | 25.321 ±<br>2.67 [28] | KW, 0.67 | ns | ns | ns |
| Max frequency (Hz) | 110.9 ±<br>28.1 [15] | 131.2 ±<br>25.4 [11] | 80.1 ± 7.3<br>[28] | KW, 0.19 | ns | ns | ns |
| ISI CV@ max firing | 0.19 ±<br>0.03 [14] | 0.47 ±<br>0.16 [11] | 0.19 ±<br>0.02 [27] | KW, 0.20 | ns | ns | ns |
| ISI CV2 @ max firing | 0.10 ±<br>0.13 [14] | 0.27 ±<br>0.10 [11] | 0.11 ±<br>0.01 [27] | KW, 0.33 | ns | ns | ns |

### Higher action potential threshold and more distal axon initial segment in epITCs than in ML neurons

Next, we investigated AP threshold and waveform by injecting 10ms of depolarizing current steps of increasing amplitudes (Fig. 4A). In line with findings from Burton and Urban (2014), the three cell types had similar action potential waveforms, with only the peak amplitude being significantly smaller in eplTCs (Fig. 4E, Table 1). This difference is in line with similar maximum voltages reached but higher AP threshold in eplTC than in ML’s cells (Fig. 4B-D, Table 1). Of note, when the AP threshold is normalized for capacitance, pMCs require smaller injected currents than pTCs to fire (Fig. 4B, Table 1). In contrast to the sodium channel-driven rapid depolarization and in line with molecular data (Zeppilli et al., 2021), the AP phases reliant on potassium conductance (width at half-height, WHH, afterhyperpolarization, AHP) do not differ between the three cell types, bar the higher minimum voltage reached by eplTCs (Fig. 4F-H).

**Figure 4.**
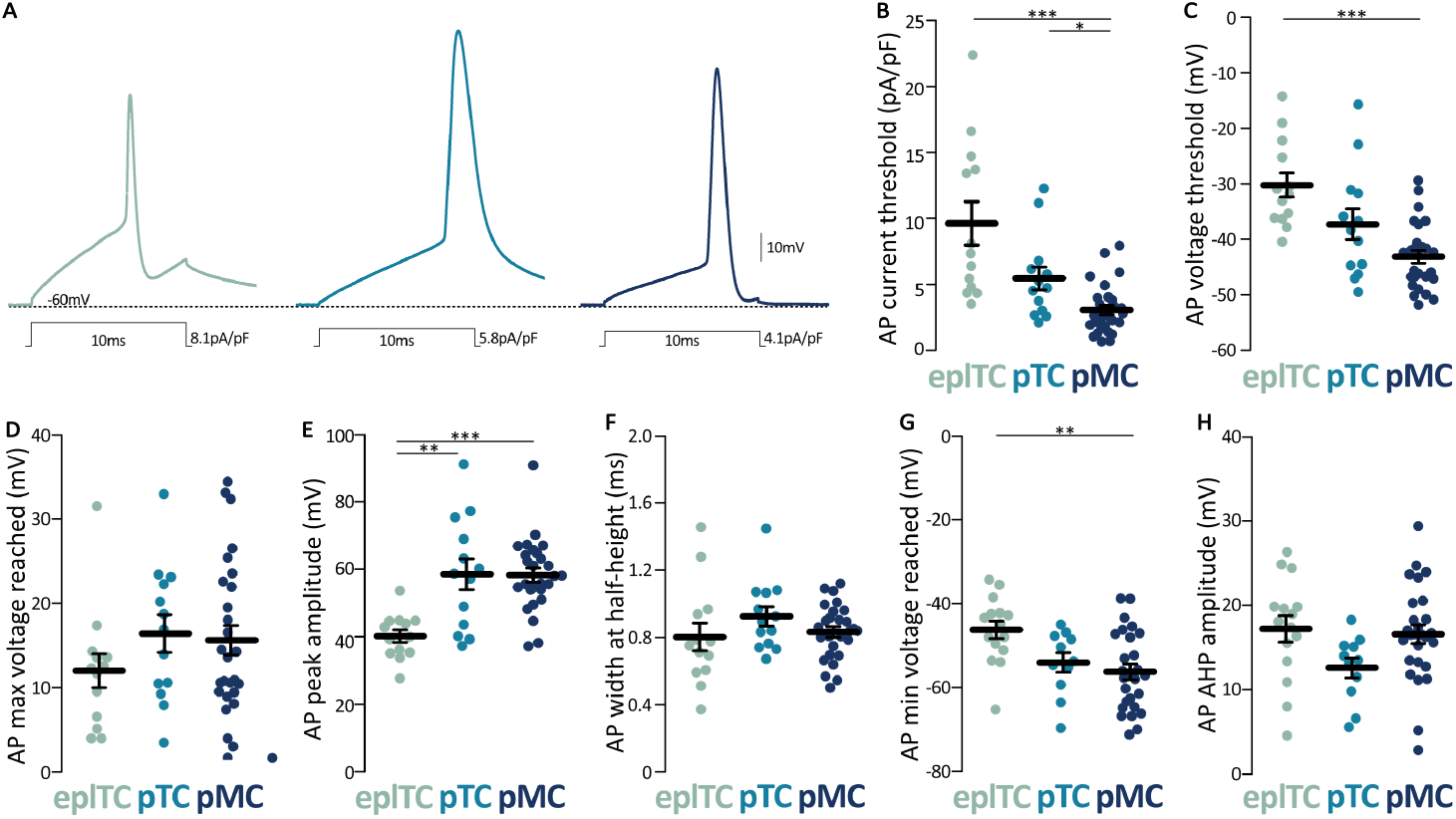
Action potential threshold and waveforms differ between OB projection neurons. **(A)** Example traces of the membrane voltage response to the minimum depolarizing 10ms-long current injection needed to evoke an action potential (AP) in epITCs (green, n=13), pTCs (cyan, n=13) and pMCs (blue, n=27). Waveform parameters for the three OB projection neurons subtypes include **(B)** Injected current density needed to evoke an AP, **(C)** membrane potential at which the AP was evoked, **(D)** maximum voltage reached by the AP, (E) AP peak amplitude, **(F)** AP width at half the maximum height, **(G)** maximum voltage reached by the AP, and **(H)** peak amplitude of the AP after hyperpolarization (AHP). Circles are individual cells, lines are mean ± SEM, *p<0.05, **p<0.01, ***p<0.001; further quantification and statistical analysis in Table 1.

In summary, despite being more electronically compact, eplTCs require more current to fire an action potential which however has a similar shape to ML cell APs.

To investigate whether this threshold difference may be due to morphological differences at the AP initiation site, *i*.*e*. the axon initial segment (AIS), we performed immunohistochemistry against the AIS master organiser and label ankyrinG (AnkG; Fig. 5A)(Kole et al., 2007; Bender and Trussell, 2012; Leterrier, 2018; Galliano et al., 2021). We found no difference in AIS length between the three cell types (eplTC = 23.28 ±1.41 µm; pTC= 23.28±0.80 µm; pMC = 25.07±0.72 µm; ANOVA nested on mouse F(2, 11)=0.5529, p=0.59; Fig. 5B and D). Conversely, eplTCs’ AISes are extremely distal compared to those of principal neurons in the ML (eplTC = 21.19±4.78 µm; pTC = 2.88±0.25 µm; pMC= 5.02±0.46 µm; log-transformed data for ANOVA nested on mouse = F(2,11)=14.16, p*<*0.001, post-hoc with false discovery rate correction, eplTC vs pTC p*<*0.001; eplTC vs pMC p=0.003, pTC vs pMC p=0.13; Fig. 5C).

**Figure 5.**
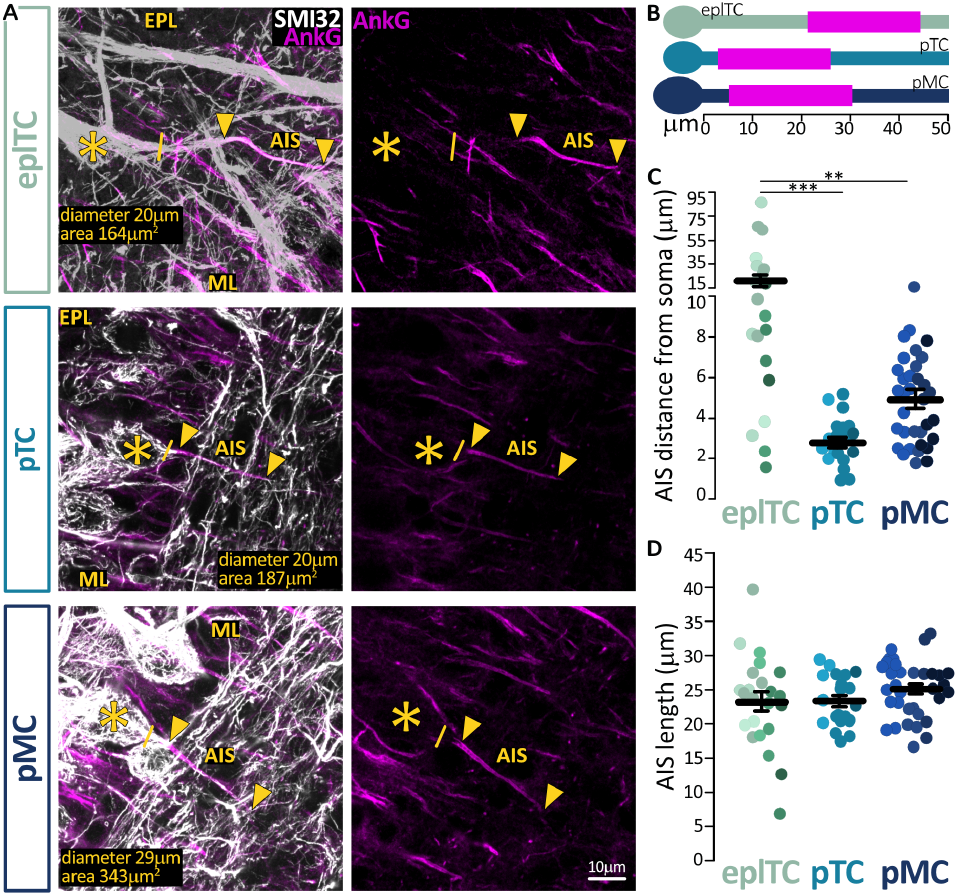
EPL cells have similarly long but more distal axon initial segments than ML neurons. **(A)** Example maximum intensity projection images of epITC, pTC and pMC neurons visualized via SMI-32 immunolabel (white) with an identified AnkG-positive AIS (magenta, arrows). Solid line indicates the emergence of the axonal process from the soma (asterisk). EPL = external plexiform layer, ML = mitral layer. **(B)** Mean AIS start and end position for each group. **(C-D)** AIS distance from soma and length in epITCs (green, n=23), pTCs (cyan, n=21) and pMCs (blue, n=34). Circles represent individual cells, different colors represent different mice, lines are mean ± SEM, **p<0.01, ***p<0.001.

Recent work has identified axons originating from dendrites not just in inhibitory bulbar interneurons but also in pyramidal cells in cortex (Galliano et al., 2021; Hodapp et al., 2022), and such dendritic origin can correlate with a more distal AIS location. However, we confirmed that while 8/23 eplTCs had dendritic axons, their AISes were not different from the AISes of the 15/23 eplTCs in terms of distance from soma, length, or diameter (nested t-tests, all p*>*0.09). Importantly, the diameter of both the proximal axon and AIS were identical not just between eplTCs with somatic or dendritic-origin AISes, but among all cell types (ANOVA nested on mouse, F(2, 10)=1.28, p=0.32, F(2, 10)=1.47, p=0.28 respectively). Literature suggests that, at comparable diameters and lengths (Goethals and Brette, 2020), excitability reduces the further the AIS is from the soma as more charge is required to overcome charge dissipation and generate an AP (Yamada and Kuba, 2016). Thus, this morphological data could at least partially explain the higher firing threshold in eplTC, but fails to account for the difference in firing threshold recorded between pMCs and pTCs in the mitral layer.

### Repetitive action potential properties are comparable across putative cell types

To investigate rate and temporal coding in projection neurons, we injected longer current steps of increasing intensity (Fig. 6A-D). To account for different cell capacitance, we constructed input-output curves with injected current density as the independent variable (two-way ANOVA, effect of cell type F(27-220)=27.08, p*<*0.001; effect of current density F(14-220)=2.24, p=0.07; effect of interaction F(28-220)=2.06, p=0.002; Fig. 6E). From them, we extracted rheobase (Fig. 6F) and the rate of rise by fitting a line and calculating its slope (Fig. 6G).

**Figure 6.**
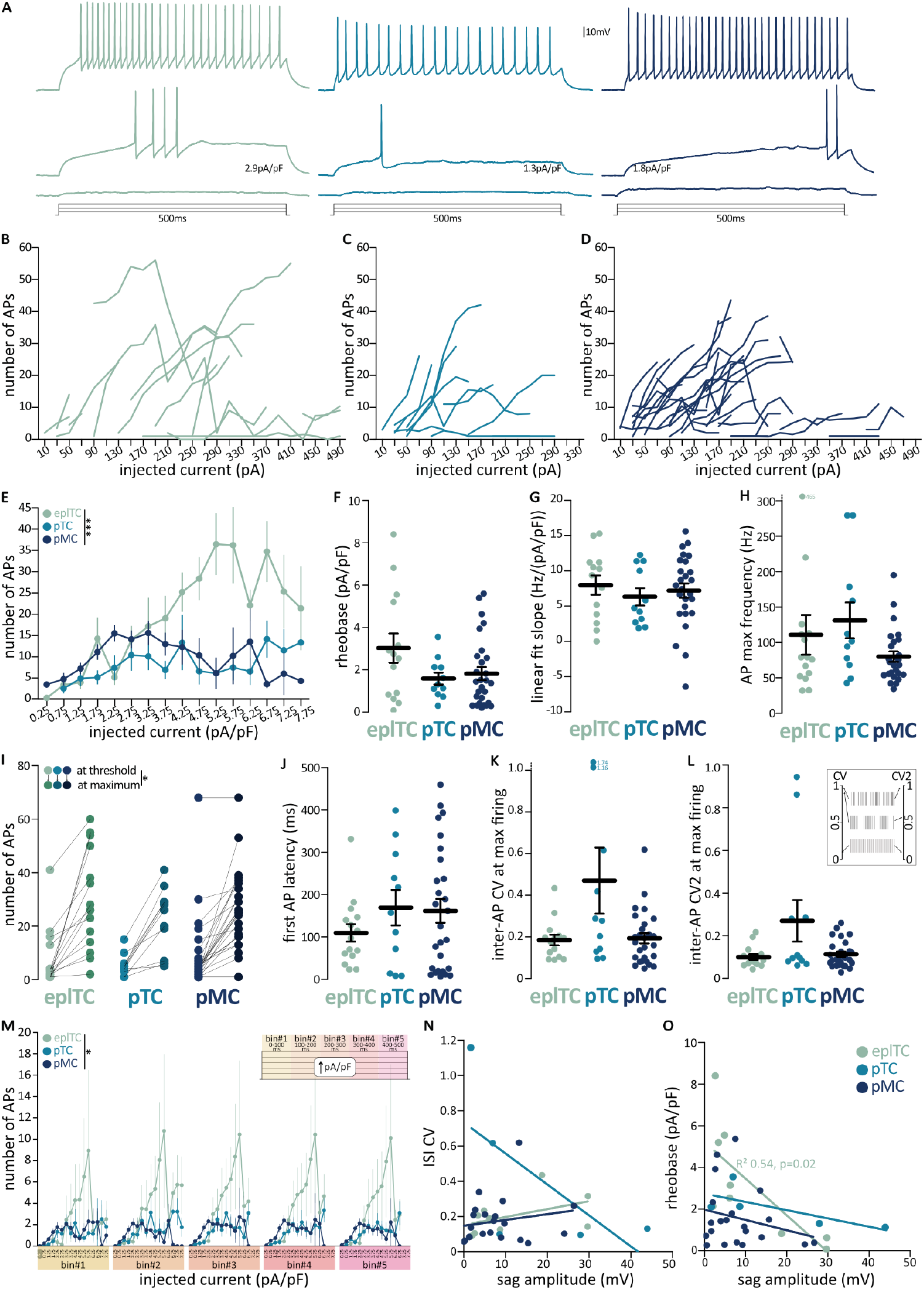
Comparable repetitive action potential firing among OB projection neurons. **(A)** Example traces of the membrane voltage response to 500ms depolarizing current injections at 300pA, rheobase, and max AP firing in epITCs (green, n=l5), pTCs (cyan, n=11), and pMCs (blue, n=27). **(B-D)** Input-output plots showing the number of APs fired at each current injection in individual neurons. **(E)** Mean number of APs and SEM at each current density *(i*.*e*., injected current normalized for cell capacitance) in epITCs (green), pTCs (cyan), and pMCs (blue). Repetitive firing parameters include **(F)** rheobase, **(G)** slope of these number of APs vs. current density input-output curves, **(H)** maximum AP firing frequency, **(I)** number of APs at threshold (light shades) and at maximum firing (dark shades), **(J)** Latency of the first AP at the current injection level where the max AP number was fired, **(I-J)** coefficient of variation of the inter-spike intervals (ISIs, CV) and of adjacent ISIs (CV2; see inset for graphical description) at the current injection level where the max AP number was fired. **(M)** Input-output curve with the 500ms current injection divided into 5 100ms bins (see insert). **(N-O)** Correlation of sag amplitude with firing properties. Circles and thin lines are individual cells, thick black lines are mean ± SEM, *p<0.05, ***p<0.001; further quantification and statistical analysis in Table 1.

In line with the current and voltage threshold results discussed above, also with these longer-lasting injections eplTC took longer than ML cells to fire APs, the rheobase values did not reach significance (Kruskal Wallis test, p=0.08), and all three cell types had similarly steep input-output relationships (Table 1). We also found no difference between the three cell types in the latency to fire the first AP, in the number of APs threshold or at maximum firing, nor in the maximum AP frequency (Fig. 6H-J, Table 1). Both *in vivo* and *ex vivo* preparations have shown that TCs fire more irregularly than MCs (Burton and Urban, 2014; Fourcaud-Trocmé et al., 2022).

To investigate firing regularity, we calculated both the inter-spike intervals (ISI) coefficient of variation (CV - where a high ISI CV indicates irregular firing, Fig. 6K) and the coefficient of variation of adjacent intervals (CV2, sensitive to regularity within a burst, Fig. 6L). Surprisingly, we found no differences between the three cell types in each measure, both at maximum firing (Table 1) and rheobase (data not shown). We analysed temporal coding further by binning the current injection into five 100ms epochs and plotted the firing along them (Goldfarb et al., 2007). We found that while eplTCs are more excitable, all three cell types fire similar number of action potentials at the beginning, middle, and end of the current injection (bin average AP number two-way ANOVA, effect of cell type F(2,125)=3.58, p=0.03; effect of current density F(4,125)=0.08, p=0.99; effect of interaction F(8,125)=0.05, p=1.00; Fig. 6M). Finally, we checked if the sag voltage amplitude correlated with either firing CV or rheobase (Burton and Urban, 2014), but found a significant correlation only for sag amplitude and rheobase in eplTCs (linear regression R2=054, F=8.22, p=0.02; all other correlations p*>*0.16; Fig 6O).

Taken together, our results indicate that when capacitance is accounted for, eplTCs have similar firing patterns to both pTCs and pMCs in the mitral layer.

### Can putative cell type be accurately classified into discrete groups?

Despite the wealth of literature suggesting that TCs and MCs are physiologically different, our *ex vivo* data indicate that individual physiological passive and active properties are not sufficient to differentiate them. If individual properties cannot return a clean linear classifier, does the mitral vs tufted split appear when all these values are considered holistically? To answer this question and determine the source of the variation in our dataset, we performed a PCA including all measurements extracted from passive properties and AP firing protocols (Fig. 7A). The first three principal components cumulatively accounted for 50% of the variance and the most influential loading scores were connected to AP amplitude and threshold, as well as capacitance (Fig.7B-C). While on average eplTCs and pMCs clustered at opposite ends of the PC1 axis and pTCs were more narrowly grouped around the middle, we found considerable overlap between the three cell types.

**Figure 7.**
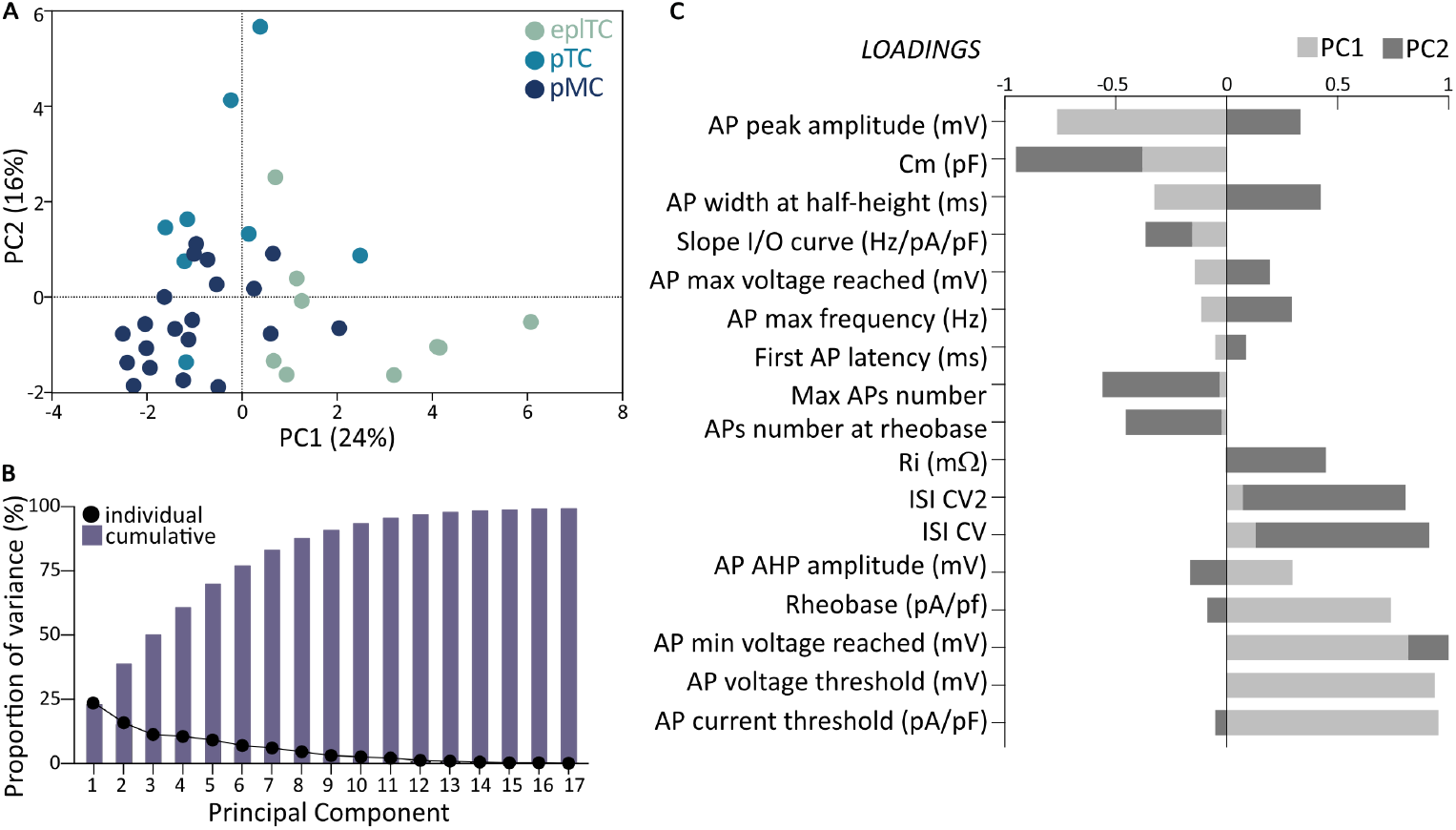
Principal component analysis (PCA) of firing properties fails to reveals clear clustering of OB projection neurons. **(A)** PC score plot for epITC (green, n=9), and mitral layers’ pTC (cyan, n=8) and pMC (blue, n=20) based on passive properties and all measurements obtained from AP firing recordings (Figures 2-4-6, Table 1). Each circle represents a cell plotted against its primary and secondary principal component (PC) scores. **(B)** Individual (circles) and cumulative (bars) proportion of variance explained by each principal component. **(C)** Loading scores for all variables showing their respective contributions to PCI (light grey) and PC2 (dark grey).

In summary, the PCA failed to return clear clustering, but suggested a more gradual continuum of heterogeneous properties in OB projection neurons.

## Discussion

In this study we compared the morphological and functional properties of bulbar principal neurons across EPL and ML using classification approaches based on agreed-upon soma size and unbiased clustering of membrane capacitance. We were unable to conclusively segregate pMCs and pTCs within the ML, but we confirmed earlier findings that very large ML cells are overall different from TCs in the EPL. Historically, smaller cells in the ML and cells with their soma only partially in the ML have been excluded from analysis because of their ambiguous identity (Burton and Urban, 2014; Nagayama et al., 2014). If these “transitional” cells are ignored, a very clear mitral-tufted split emerges. However, when they are included, such stark classification blurs as summarized by our PCA of action potential properties which returned a polarized continuum rather than defined clusters. Taken all together these data suggest that, besides the often-unavailable connectivity profile, anatomy paired with size and location is still the best classifier of bulbar principal neurons, which are overall extremely heterogeneous.

### Age, recording conditions, and analysis methods strongly influence AP firing

Both *ex vivo* and *in vivo* studies have shown that TCs cells are more excitable than MCs, and that they have a latency to fire in response to OSN inputs (Burton and Urban, 2014). While for 500ms-long current injections we observed higher numbers of APs in eplTC input/output curves, our dataset did not replicate this higher excitability finding when looking more granularly at single APs thresholds. This discrepancy is likely due to both experimental and analysis differences. Indeed, contrary to Burton and Urban (2014) we recorded at physiologically-relevant temperatures without synaptic blockers in post-weaning mice, and then normalized the injected shorter-lasting current injections for cell capacitance, which is by definition very different between the groups. Both age and synaptic blockers have been shown to be strong modulators of MC firing (Smithers et al., 2017; Van Hook, 2020; Tufo et al., 2022), which with increasing age becomes more attuned to high frequency stimuli (Yu et al., 2015) and as such can easily explain the discrepancy. Moreover, our AIS morphological data are in line with the higher firing threshold in eplTCs. The AIS location, which together with its morphology has been shown to correlate best with somatic threshold than rheobase, was very distal in eplTCs. Given the similar proximal axon diameters in eplTCs and ML cells, this suggests that eplTCs do not operate in a high coupling regime with the soma and thus need more charge to initiate firing (Brette, 2013; Goethals and Brette, 2020). Furthermore, despite variations in recording conditions, our dataset mirrored the heterogeneity in sag potentials reported previously (Angelo and Margrie, 2011; Burton and Urban, 2014), and in general in most single AP and repetitive firing parameters.

### Mitral, tufted, and everything in between: grey areas in the classification of bulbar principal neurons

The cell-to-cell variability within and between putative subclasses that we report here stresses that bulbar principal neurons are a heterogenous population (Zeppilli et al., 2021). Including both the analysis of soma size in fixed tissue and the firing properties, our dataset failed to return a clear separation between pMCs and pTCs in the ML, and even a ML vs EPL classification is somewhat blurred. This is not surprising because, while a traditional location + 20µm-diameter classifier is appealing, there is accumulating evidence of overlap in the molecular and genetic profiles of MCs and TCs. For example, they heterogeneously express GABA_A_ receptors and voltage gated potassium channels (Panzanelli et al., 2005) and while MCs and TCs can be classified at the level of transcriptomics, cell-type-specific modules of gene regulation fail to granularly class MCs and TCs (Zeppilli et al. 2021). Indeed, it is thought that their biophysical diversity is at least partly due not to genetic programs, but to experience dependent factors which have the potential to expand their heterogeneity (Padmanabhan and Urban, 2010; Angelo et al., 2012; Tripathy et al., 2013).

Importantly, OB principal neurons also differ in their morphology of lateral dendrites and their axonal projections, and have consequent differences in synaptic connectivity (Christie et al., 2001; Nagayama, 2010; Gire et al., 2012; Nagayama et al., 2014; Geramita et al., 2016; Liu et al., 2019; Imamura et al., 2020; Jones et al., 2020). The combination of structural, functional, intrinsic and synaptic properties likely returns not two clean groups – MCs and TC – but multiple subgroups (Padmanabhan and Urban, 2010; Angelo and Margrie, 2011; Kikuta et al., 2013; Goaillard and Marder, 2021; Zeppilli et al., 2021). This broad diversity could enable the olfactory bulb to parallelly represent the wide odour and concentration space (Lee et al., 2023) via spatially and temporally distributed ensembles of active neurons (Uchida et al., 2014; Geramita et al., 2016; Shmuel et al., 2019).

### Heterogeneity as a key odour processing tool

For most sensory systems the stimulus space is relatively well known (Huberman and Niell, 2011; King et al., 2015; O’Connor et al., 2021). Conversely we still do not know how many odours a rodent – or human – could potentially detect, but we know that the range is truly vast (Meister, 2015; Lee et al., 2023) and that it does so via a regenerating peripheral sensor (Schwob, 2002). Olfactory processing is further unique because it eschews a thalamic relay and information only takes two synapses to go from nose to cortex and other higher areas (Shepherd, 2004). Given this stimulus and sensor complexity and such paired down relay anatomy, it is not surprising that the OB needs more than two parallel channels to process odours. It is thus tempting to speculate that heterogeneity of genes, morphologies, intrinsic properties, and synaptic connections is used throughout the olfactory system – principal neurons as well as OSNs and interneurons (Godfrey et al., 2004; Antal et al., 2006; Lledo et al., 2008; Galliano et al., 2018) - to sense, process, and classify olfactory information.

## Author Contributions

SG: Investigation, Validation, Formal analysis, Data curation, Visualization, Writing – original draft, Writing – review and editing

CG: Investigation, Writing – review and editing LH: Investigation, Formal analysis

SJBW: Validation, Software NB: Formal analysis

PHP: Funding acquisition, Supervision, Writing – review and editing

EG: Conceptualization, Investigation, Data curation, Formal analysis, Visualization, Funding acquisition, Project administration, Supervision, Writing – original draft, Writing – review and editing

The authors declare no financial or personal relationship which could be construed as a potential conflict of interest.

## Acknowledgments

This work was supported by a UKRI Medical Research Council Equipment Grant (MC-PC-MR-X012271/1) and project grants from the Royal Society (RGS-R1-19148), the URKI Biotechnology and Biological Sciences Research Council (BB-W014688-1) and the Newton Trust (EG); an Icelandic Research Fund Project Grant 217945-051 (PHP,EG); a Cambridge Trust PhD studentship (LH); and a University of Cambridge Institute of Neuroscience postgraduate scholarship (SJBW). We wish to thank Sue Jones, Matthew Grubb and Ailie McWhinnie for comments on the manuscript, and all members of the Galliano laboratory for providing helpful discussions.

